# Negative regulation of p53 by the poliovirus receptor PVR is a target of a human cytomegalovirus immune evasion molecule

**DOI:** 10.1101/2022.07.04.498680

**Authors:** Adam F. Odell, Aarren J. Mannion, Pamela F. Jones, Graham P. Cook

## Abstract

Initially characterised for its role in maintaining genome integrity, p53 has emerged as a critical hub for coordinating cellular responses to diverse types of stress. Here we identify cell surface receptor loss as a signal for p53 induction. The poliovirus receptor (PVR) regulates angiogenesis, leucocyte adhesion and immune surveillance. We demonstrate that loss of PVR from endothelial cells also promotes cell cycle arrest through the induction of a p53 transcriptional programme. The p53 induction is post-translational and, despite remaining associated with MDM2, p53 exhibits reduced ubiquitination, aiding its stabilisation. Increased expression of PVR marks malignant or infected cells, and retention of PVR in the endoplasmic reticulum by human cytomegalovirus (HCMV) UL141 protein allows HCMV infected cells to evade immunity. We show that intracellular retention of PVR by UL141 prevents p53 induction, allowing HCMV to escape both the immune- and p53-mediated surveillance functions of PVR. These data reveal that p53 coordinates responses to changes in cell surface composition and that the cell intrinsic PVR-p53 pathway coupled with PVR-mediated immune surveillance functions provide a sensor mechanism to maintain expression of this multi-functional cell surface molecule.

## Introduction

The p53 molecule is a transcription factor with tumour suppressor activity stemming from its ability to negatively regulate cell growth and proliferation (1–3). The first signals shown to induce p53 activity were those resulting from DNA damage. Genotoxic stress, such as that induced by X-rays, chemical carcinogens and chemotherapeutic agents, initiates a DNA damage response that results in ATM/ATR dependent activation of CHK1/CHK2 kinases which then phosphorylate p53 (4, 5). This p53 phosphorylation disrupts association with the ubiquitin ligase MDM2, reducing p53 ubiquitination and limiting its degradation by the proteasome (6, 7). Increased stability allows the accumulation of p53 in the nucleus and the regulation of target gene expression (8, 9). Amongst its many transcriptional targets, p53 induces expression of the cyclin dependent kinase (CDK) inhibitor p21, arresting cells in the G1 phase of the cell cycle (10, 11); p53 also stimulates expression of proapoptotic BAX as well as MDM2, the latter providing negative feedback via increased p53 degradation (12, 13). Overall, this activity prevents the propagation of cells harbouring mutations and it is the inactivation of p53, either by mutation or by viral oncogene products, that highlight the key role that p53 responses play in tumour progression and cell fate decisions (14–17). The induction of p53 transcriptional activity in response to DNA damage led to p53 being referred to as the “guardian of the genome” (1). It has since emerged that p53 induction is not confined to DNA damage responses, but can occur as a result of other, diverse types of cellular stress including death receptor activation, hypoxia, reactive oxygen species, oncogene signalling, oxidative stress and limiting supplies of key metabolites (2, 3). These diverse signals lead to p53 stabilisation, cell cycle arrest and ultimately a spectrum of outcomes, ranging from repair and recovery, through cellular senescence, to cell death (15).

The poliovirus receptor (PVR, also known as CD155 or Necl-5) is a type I transmembrane immunoglobulin superfamily member, first identified as a broadly expressed receptor for poliovirus (18). Subsequent studies showed that PVR mediates interactions between leucocytes and endothelial cells (EC) during extravasation; EC PVR molecules interact with the leucocyte receptor DNAM-1 (also known as CD226) to facilitate leucocyte binding at EC junctions, leading to transendothelial migration (19). The interaction between PVR and DNAM-1 was discovered whilst searching for ligands of natural killer (NK) cell receptors; two DNAM-1 ligands were identified, nectin-2 (CD112) and the nectin-like family molecule PVR (20, 21). Expression of these molecules by target cells results in engagement of the NK cell activation receptor DNAM-1, triggering target cell killing and pro-inflammatory cytokine release (22, 23). Subsequently, PVR was shown to interact with two additional receptors expressed by NK cells and T cells; CD96 (originally named T cell activation, increased late expression or TACTILE) and T cell immunoreceptor with Ig and ITIM domains (TIGIT). These additional receptors compete efficiently with DNAM-1 for PVR, but deliver negative signals upon ligation, thereby acting as immune checkpoints (23). The PVR molecule can therefore regulate immune responses both positively and negatively.

The induction of PVR expression is linked to the RAS-MAPK pathway, to DNA damage via the ATM/ATR pathway and to Toll-like receptor (TLR) ligation, thus providing a mechanism to link immune surveillance to malignancy and infection (24–27). The importance of PVR in NK cell responses is best demonstrated within the context of viral infection, as both human cytomegalovirus (HCMV) and HIV reduce cell surface PVR expression to evade detection (28, 29). In HCMV, the unique long (UL)141 gene product prevents the cell surface expression of both PVR and nectin-2, thereby allowing infected cells to evade detection by DNAM-1 expressing NK cells (28, 30). Here we show that loss of PVR expression triggers a quiescent-like phenotype via a p53 dependent, cell intrinsic pathway. This identifies a new role for PVR and establishes p53 as a hub to coordinate cell fate decisions dictated by changes in cell surface composition.

## Results

### Endothelial cell proliferation, migration and angiogenic activity is regulated by PVR

The PVR molecule is widely expressed (18) and endothelial cells (ECs) have particularly high expression compared to other cell types examined (Fig. 1*A*). PVR is readily detected at the EC surface, where it is primarily localised at cell borders (Fig. 1*B* and Supplementary Fig. *1A* and *B*). Upon cell-cell contact, PVR is endocytosed, leading to reduced growth factor signalling and contact inhibition (31–33). Comparing confluent and sub-confluent cultures revealed a trend towards higher PVR expression in sub-confluent cells (Supplementary Fig. *1A-D*), as previously shown using mouse fibroblasts (31). These results suggested a link between PVR expression and EC growth. Further investigation showed that higher PVR levels in sub-confluent primary ECs were associated with higher levels of expression of several growth and proliferation markers (Supplementary Fig. 1*D*). This association was not evident in the SV40/hTERT immortalised brain EC line, hCMEC/D3 (Supplementary Fig. 1*D*), suggesting that the link between PVR expression and EC growth is disrupted in this cell line in which normal cell cycle control has been subverted by immortalisation (34).

**Fig. 1.**
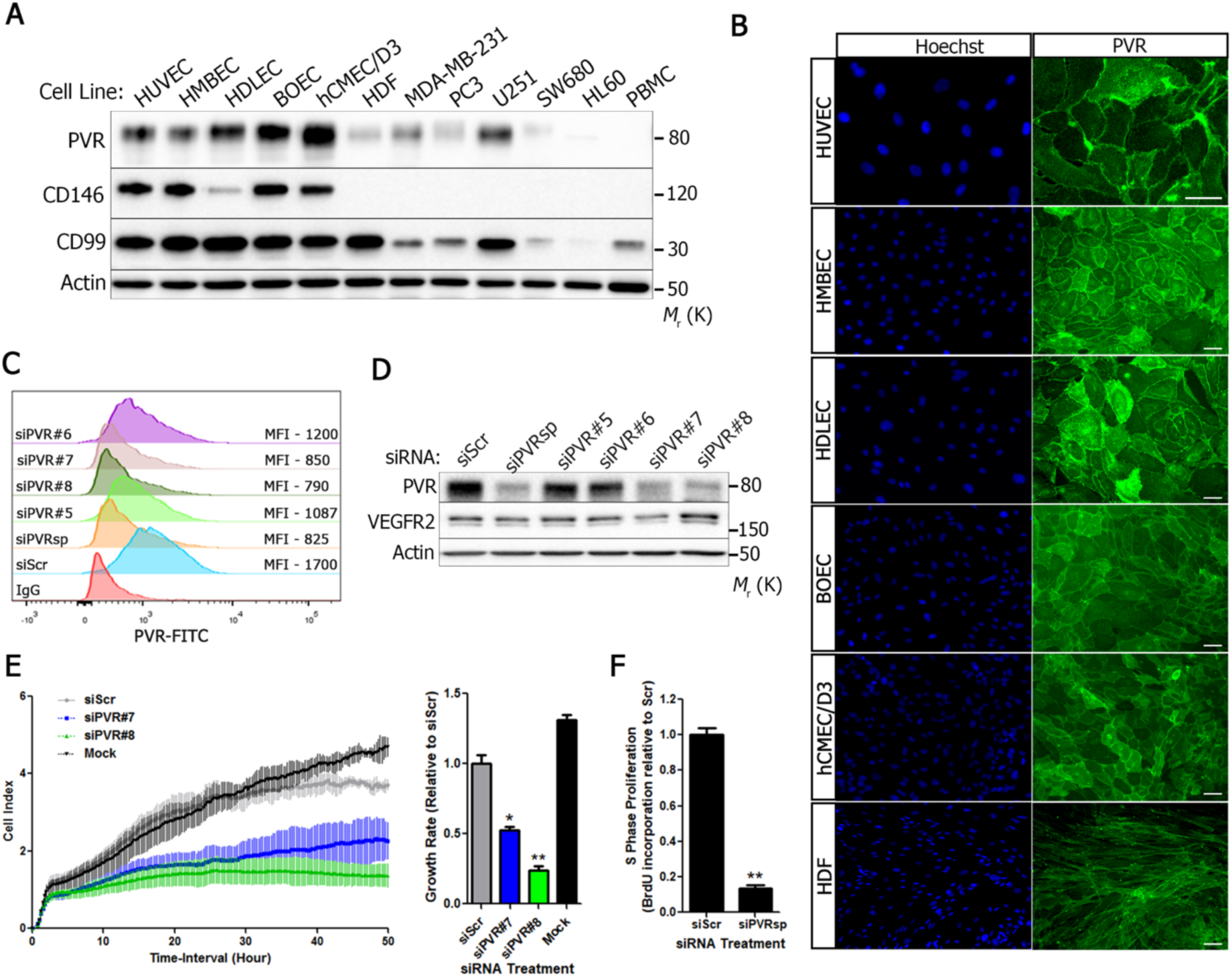
PVR expression regulates endothelial cell function. **A)** Primary ECs and a range of cell lines analysed by immunoblotting for total PVR, CD146, CD99 and actin. The primary cells are; human umbilical vein endothelial cells (HUVEC), human microvascular brain endothelial cells (HMBEC), human dermal lymphatic endothelial cells (HDLEC), blood outgrowth endothelial cells (BOEC), human dermal fibroblasts (HDF) and peripheral blood mononuclear cells (PBMC); cell lines used are hCMEC/D3, an immortalised EC line (34) and the tumour-derived cell lines, MDA-MB-231 (breast), PC3 (prostate), U251 (brain), SW680 (colon) and HL60 (monocytic leukaemia). **B)** Expression of PVR analysed by immunofluorescence microscopy on ECs and primary human dermal fibroblasts (HDFs); scale bar, 20 µm. **C)** HUVECs transfected with siRNA against PVR (or controls) for 72 h before analysis of surface PVR expression by flow cytometry; smart pool, siPVRsp; individual siRNAs for PVR, siPVR#n; scrambled control, siScr. The mean fluorescence intensity (MFI) from each histogram is indicated. **D)** HUVECs transfected with siRNA as in (*C*), analysed for PVR and VEGFR2 expression by immunoblotting. **E)** Growth of siRNA transfected HDLECs, cultured on E-plates in an xCELLigence Real Time Cell Analyser (RTCA) for 48h. Growth rates (from left hand panel) were expressed as growth rate relative to siScr transfected cells (right-hand panel). Significance was determined by ANOVA where * p<0.05 from 3 independent experiments, where each treatment condition was performed in duplicate. **F)** S phase proliferation of siRNA transfected HDLECs (measured by BrdU incorporation and ELISA) expressed relative to siScr-treated cells. Significance determined by one-tail t-test from 3 independent experiments performed in triplicate. *** p<0.001.

We used RNA interference to investigate EC PVR function in more detail, using a pool of siRNAs targeting PVR (siPVRsp) and four individual siRNA molecules from this pool (siPVR#5-8). Flow cytometry and immunoblotting showed that siPVRsp, siPVR#7 and siPVR#8 showed efficient reduction in expression of cell surface and total PVR (Fig. 1*C-D* and Supplementary Figure 2*A-C*). We used these siRNAs to investigate the link between PVR expression and EC growth. Treatment of ECs with either siPVR#7 or siPVR#8 significantly reduced growth over 48 hours (Fig. 1*E*) and PVR loss was accompanied by a reduction in the proportion of cells entering S phase (Fig. 1*F*). As expected, both siRNA and antibodies targeting EC PVR reduced the binding of EC to peripheral blood mononuclear cells (Supplementary Fig. 2*D* and *E*), consistent with the role of EC PVR in leucocyte adhesion and transmigration (19, 35). In addition, loss of PVR reduced EC migratory capacity (Supplementary Fig. 2*F* and *G*). Both proliferation and migration are key processes in angiogenesis (36) and both the collagen overlay assay (37) and fibroblast-EC co-cultures (38) showed that PVR reduction inhibited the ability of ECs to form tubule networks *in vitro* (Supplementary Fig. 2*H-J*). These results show that reduction in EC PVR expression induces a cell cycle arrest and reduced migratory activity, culminating in reduced angiogenic potential.

### Loss of PVR induces the p53 pathway in endothelial cells

Previous studies have demonstrated that p53 induction inhibits EC migration and angiogenic activity (39). The related phenotype observed here suggested a link between EC PVR loss and p53 induction. In HUVECs, a reduction in PVR by either siRNA or shRNA, induced p53 protein expression and the products of two transcriptional targets of p53, the ubiquitin ligase MDM2 (a negative regulator of p53 stability) (6, 7, 12, 40) and the CDK inhibitor, p21/CDKN1A (10) (Fig. 2*A* and Supplementary Fig. 3*A and B*). However, expression of the ubiquitin-specific protease USP7/HAUSP, which stabilises p53 and causes growth arrest (41), was unaffected by PVR depletion (Fig. 2*A*). Immunofluorescence microscopy of siPVR transfected HUVECs demonstrated an inverse relationship between cell surface PVR and nuclear accumulation of p53 (Fig. 2*B*). Indeed, transient siRNA transfection allowed the identification of ECs that lacked cell surface PVR but had readily detectable nuclear p53 (Fig. 2*B*; open arrows) alongside ECs which retained PVR, but lacked detectable p53 (Fig. 2*B*; closed arrows). The loss of PVR from ECs was detected within 24hrs of siPVR transfection and induced a similarly rapid induction of p53 and p21, which was sustained over a 72h period (Fig. 2*C*). We extended this analysis to a panel of primary ECs and the SV40/hTERT immortalised hCMEC/D3 cell line; the induction of p53, MDM2 and p21 upon PVR loss was common across all of the primary EC cultures (Fig. 2*D*). Additionally, upon PVR loss, we identified induction of BAX (also a transcriptional target of p53) (13) and reduced expression of positive regulators of the cell cycle, namely cyclin A2, cyclin B1, CDK1, CDK2 and PCNA in the primary ECs (Fig. 2*D*). However, the hCMEC/D3 cell line constitutively expresses high levels of p53 which were not further enhanced following PVR knockdown and hCMEC/D3 cell cycle regulators were largely unaffected (Fig. 2*D*).

**Fig. 2.**
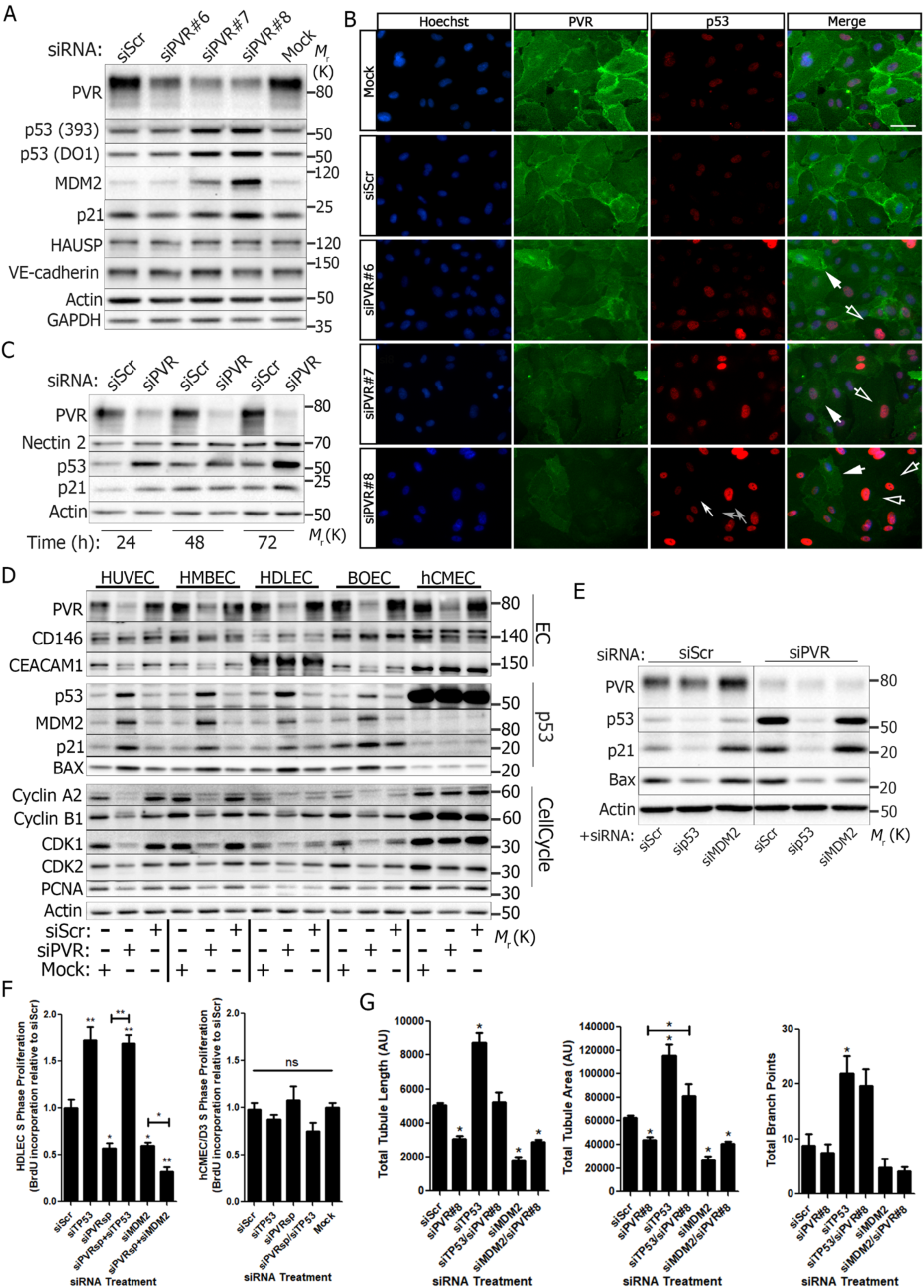
PVR depletion induces nuclear p53 accumulation, cell cycle arrest, and impaired angiogenesis. **A)** HUVECs transfected with siRNA against PVR for 72 h were examined for activation of p53 pathway components (as indicated) by immunoblotting. **B)** Immunofluorescence microscopy of HUVECs analysed for intracellular p53 and cell surface PVR expression following PVR depletion with indicated siRNA, via transient transfection. Filled arrows indicate cells where PVR expression is maintained and nuclear p53 is suppressed. Unfilled arrows highlight cells with absent PVR exhibiting increased nuclear p53. Scale bar: 20 µm. **C)** PVR and p53 expression examined 24 h, 48 h and 72 h after PVR siRNA transfection of BOECs by immunoblotting. **D)** Various endothelial cell types were transfected with siRNA against PVR for 72 h prior to immunoblotting of indicated endothelial markers, p53 pathway components, and cell cycle regulators. **E)** BOECs were transfected with combinations of siRNA targeting PVR, p53 and MDM2 as shown and analysed for expression of p53 and downstream targets by immunoblotting 72 h post siRNA transfection. **F)** S phase proliferation (via BrdU incorporation and ELISA) in HDLECs (left panel) and hCMEC/D3 (right panel) treated with siRNA combinations as indicated. Transfected cells were grown for 48 h prior to reseeding at 5×10^3^/well for a further 24 h before addition of BrdU for 4 h, followed fixation and analysis of BrdU incorporation by ELISA. Significance was determined by ANOVA where * p<0.05 and ** p<0.01 from n of 3 independent experiments. **G)** Tubulogenesis in ECs treated with combinations of siRNA targeting PVR, p53, and MDM2 (or controls) as indicated. 9-day fibroblast/EC co-cultures were used to assess Total tubule length, total tubule area and average tubule length as indicated. Significance was determined by ANOVA where * p<0.05 from 3 independent experiments.

To determine whether p53 induction was critical for the cell cycle arrest program we depleted PVR from ECs, along with p53, or its negative regulator, MDM2, using siRNA; co-depletion of PVR and p53 prevented the induction of p21 and BAX that was evident upon treatment with siPVR alone (Fig. 2*E*) and prevented the nuclear accumulation of p21 (Supplementary Fig. 3*C*). Depletion of MDM2 alone also led to an increase in p21 expression, which was not substantially enhanced by PVR siRNA. Furthermore, co-depletion of MDM2 and PVR did not increase p53 levels above those induced by PVR depletion alone, identifying PVR loss as a potent signal for p53 induction, possibly by disrupting the action of MDM2 (Fig. 2*E*). Importantly, reducing p53 and PVR together restored the ability of ECs to proliferate in the absence of PVR, as determined by BrdU incorporation (Fig. 2*F*). In contrast, hCMEC/D3 cells, which fail to modulate expression of MDM2, BAX and other cell cycle regulators following PVR loss (Fig. 2*D*), do not undergo cell cycle arrest after PVR depletion (Fig. 2*F*).

There was also a pronounced recovery in the angiogenic potential of ECs depleted of both PVR and p53 when compared to PVR-depleted cells alone (Fig. 2*G* and Supplementary Fig. 3*D*). This recovery was not evident when MDM2 and PVR were co-depleted, and tubule formation was markedly reduced under these conditions (Fig. 2*G*), consistent with the inhibition of angiogenesis following p53 induction. The ability of EC spheroids to sprout when embedded in a collagen matrix was also restored upon co-depletion of both PVR and p53 (Supplementary Fig. 2*E*). Depletion of p53 together with PVR also restored normal endothelial morphology; however, depletion of MDM2 together with PVR exacerbated the defects induced by PVR loss (Supplementary Fig. 2*F*). These findings identify p53 induction as a critical step in the EC response to PVR loss and show that sustained PVR expression is required for the efficient proliferation, migration and angiogenic responses of primary human ECs.

### PVR loss stabilises p53 and induces p53 target genes

We next investigated the mechanism by which p53 accumulates following PVR reduction. The p53 molecule regulates hundreds of genes, both positively and negatively, many by direct transcriptional control (8, 42). Following siPVR treatment of HUVECs, we detected induction (*CDKN1A, MDM2, BAX, and CCND1*) and inhibition (*CDK1, CDK2, CCNA2*, and *CCNB1*), of p53 target genes (Fig. 3*A*), consistent with the protein expression data (Fig. 2*D*). However, we did not detect any change in *TP53* gene expression itself, implicating a post-translational mechanism of p53 induction. The p53 protein has a short half-life and it is highly sensitive to protein synthesis inhibition by cycloheximide (CHX). Accordingly, p53 was largely undetectable in HUVECs after 90 min of CHX treatment (Fig. 3*B*). However, pre-treatment of HUVECs with siPVR for 72hrs to elevate p53 protein levels, largely prevented p53 downregulation induced by CHX (Fig. 3*B* and *C*). Furthermore, CHX-mediated loss of MDM2 was also stabilised following siPVR treatment, reflecting increased p53 levels and enhanced transcriptional activity (Fig. 3*B*). This reveals that a pre-existing pool of p53 protein is stabilised following PVR reduction.

**Fig. 3.**
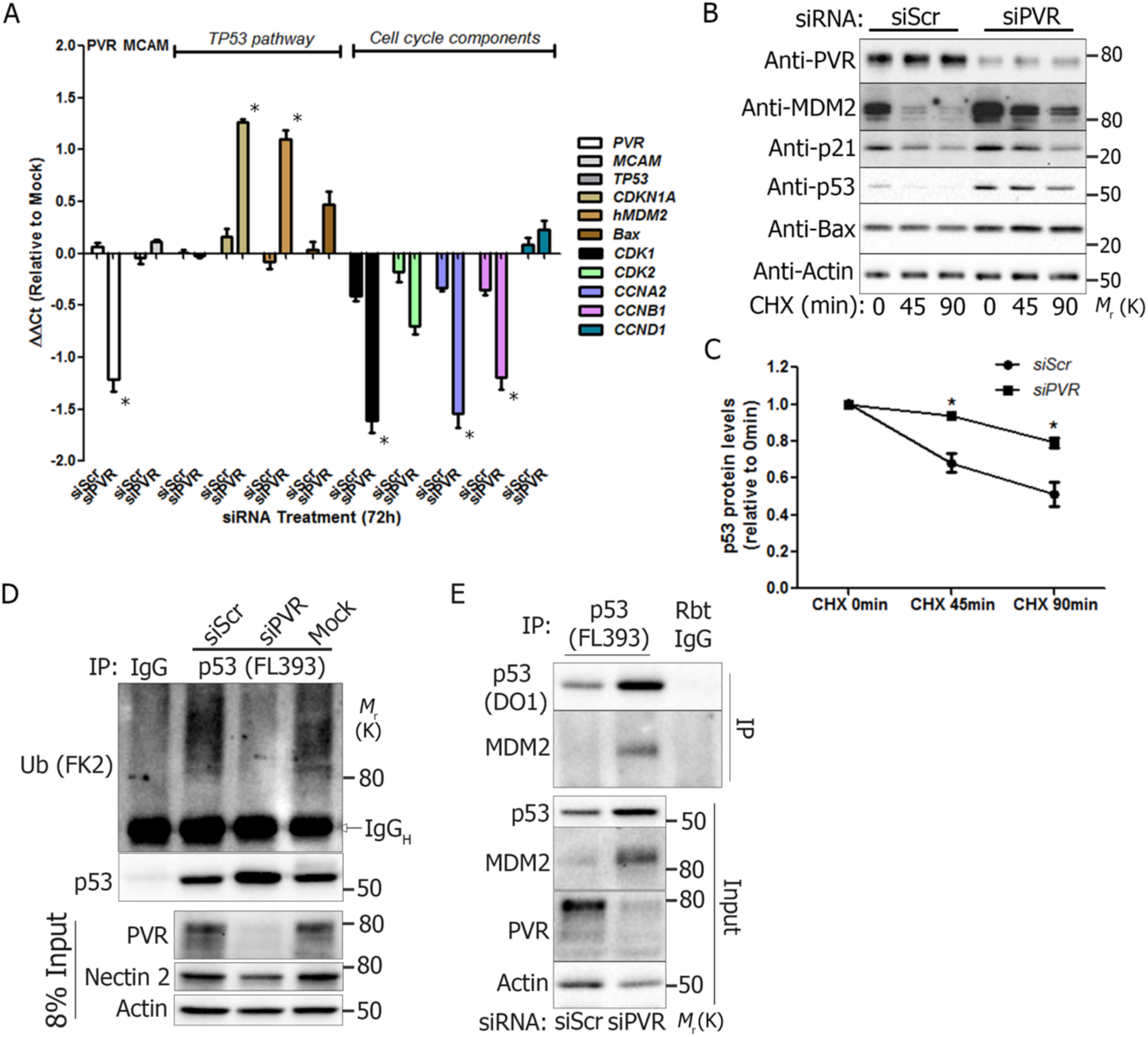
PVR controls p53 protein stability by altering p53 ubiquitination. **A)** Gene expression of p53 pathway components and downstream cell cycle regulators (determined by qRT-PCR) following transfection of ECs with indicated siRNA. Expression is reported relative to Mock treatment and standardised against GAPDH. Data represents 3 independent experiments, with each reaction performed in duplicate and analysed by ANOVA; *p<0.05. **B)** HUVECs treated with PVR siRNA were treated with cycloheximide (CHX) for 45 or 90 minutes and protein expression determined by immunoblotting with quantitation of p53 levels shown in C). **C)** p53 protein levels from B) were quantified and expressed relative to the control treatment (0hr CHX) Three independent experiments were analysed by ANOVA; *p<0.05. **D)** Total p53 was immunoprecipitated from HUVECs treated with siRNA and assayed for ubiquitin conjugation by immunoblotting using an anti-ubiquitin antibody. Input lysates were probed for PVR, Nectin-2 and actin. **E)** Anti-p53 immunoprecipitates from siRNA-treated BOECs were probed for bound MDM2 by immunoblotting.

MDM2 destabilises p53 via ubiquitin conjugation (6, 7), suggesting disruption of MDM2 E3 ligase function as a potential mechanism for p53 accumulation following PVR depletion. Indeed, treatment of HUVECs with siPVR followed by immunoprecipitation of p53 and immunoblotting with an anti-ubiquitin antibody revealed that p53 was ubiquitinated under steady state conditions and that this activity was indeed inhibited by PVR loss (Fig. 3*D*). Surprisingly, although p53 ubiquitination was reduced upon PVR depletion, co-immunoprecipitation revealed that MDM2 remained associated with p53 under these conditions (Fig. *3E*). This suggests that either MDM2 can no longer ubiquitinate p53 in the absence of PVR, or that de-ubiquitination machinery may be aberrantly activated by PVR depletion (albeit that levels of HAUSP/USP7, a major p53 DUB, were unchanged by PVR loss; Fig. 2*A*). These data show that PVR loss impairs the ability of MDM2 to mediate ubiquitination of p53, leading to p53 protein stabilisation, accumulation and induction of a p53-mediated transcriptional programme.

### Modulation of the PVR-p53 axis by a Herpesvirus immune evasion molecule

Intracellular infection and malignant transformation induce PVR expression and this favours detection and destruction by cytotoxic lymphocytes bearing DNAM-1 (20, 28). In common with other herpesviruses, human cytomegalovirus (HCMV) encodes numerous immune evasion molecules which serve to limit the detection of infected cells by both innate and adaptive immunity (43, 44). These evasion molecules include the endoplasmic reticulum (ER)-resident glycoprotein, UL141, whose expression is sufficient to block cell surface localisation of PVR and, along with HCMV Unique Short (US)2, is required to prevent nectin-2 (CD112) expression (28, 30, 45). We speculated that UL141-mediated PVR retention would facilitate immune evasion whilst retaining the PVR-p53 signalling axis, thereby maintaining key EC functions. We first analysed HCMV infection of human dermal foreskin fibroblasts (HDF). Two HDF variants, one early passage primary culture and the other, a later passage hTERT-immortalised cell line, were infected at a multiplicity of infection (MOI) of 1 or 10 with the Merlin strain of HCMV for four days. As reported previously (28, 30), HCMV infection resulted in the appearance of a lower molecular weight species of PVR, consistent with UL141-mediated retention of immature PVR in the ER (Fig. *4A*). Accumulation of this altered PVR species was associated with increased p53 and MDM2 expression, although p21 remained largely unchanged (Fig. *4A*), suggesting that the PVR-p53 axis might be disrupted. However, since HCMV infection itself induces p53 (46), we replaced HCMV with a lentivirus encoding a V5 epitope-tagged UL141 (45). Expression of UL141 resulted in the intracellular accumulation of the smaller PVR species and analysis of cell surface biotinylation profiles confirmed the intracellular retention of PVR in the presence of UL141 in HUVECs (Figure 4*B*) and fibroblasts (Supplementary Fig. 4*A*). In contrast, other plasma membrane proteins (CD146, CD99, VE-cadherin and VEGFR2) were largely unaffected (Fig. *4B*). UL141 expression in additional EC variants (HMBECs and BOECs) also demonstrated intracellular PVR retention and accumulation of lower molecular weight PVR (Supplementary Fig. *4B*). However, despite loss of PVR from the cell surface, UL141-expressing HUVECs did not accumulate p53 or MDM2 protein (Fig. 4*C*). Without p53 induction, DNA synthesis in UL141 expressing cells continued unabated, indeed, a small increase in BrdU incorporation was observed (Fig. *4D*). However, UL141 expressing ECs remained sensitive to both camptothecin (Fig. 4*D*) and nutlin-3 (Fig. *4E* and *F*), demonstrating that p53 induction in response to both DNA damage and MDM2 inhibition remained intact. Although proliferative defects were not observed, UL141-expressing HUVECs showed impairment of angiogenic functions *in vitro*, with significantly reduced network branching and average network length (Supplementary Fig. 4*C* and *D*). This reveals that cell surface PVR is required for optimal EC angiogenic function, independent of the PVR-p53 axis and the regulation of proliferative responses.

**Fig. 4.**
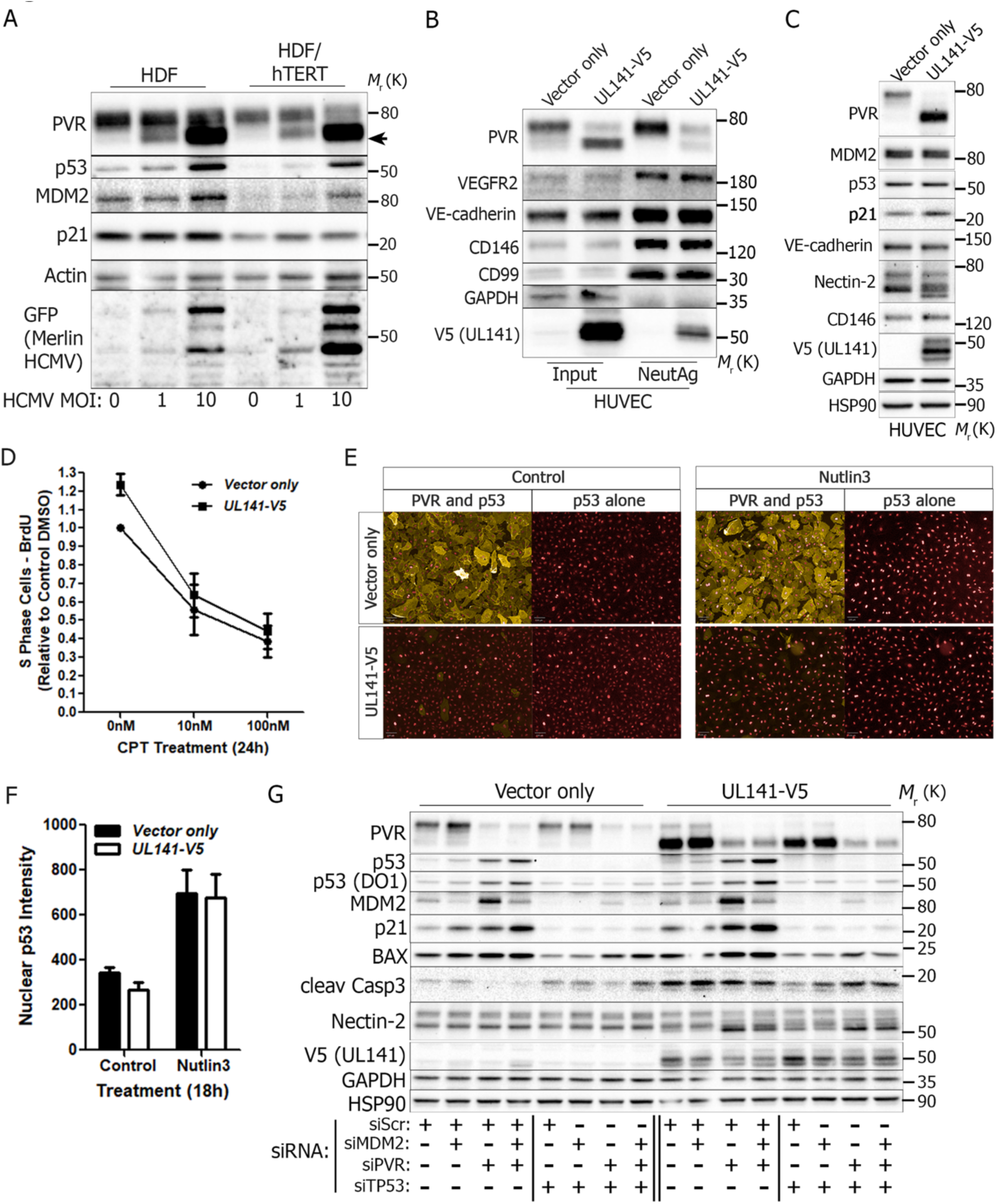
HCMV UL141 maintains PVR in the ER but does not induce p53 accumulation. **A)** Two variants of HDFs (primary and hTERT immortalised) were infected with the Merlin strain of HCMV at a Multiplicity of Infection (MOI) of 1 or 10, for 72 h prior to western blotting of the indicated proteins. **B)** Lentiviral expression of V5-tagged UL141 (pLV-UL141), or empty vector (pLV-Scr) in HUVECs and analysis of total (input) or cell surface (NeutAg) proteins following biotinylation and neutravidin-agarose pulldown. **C)** HUVECs expressing UL141 (or control) were assessed for changes in expression of the indicated molecules using western blotting. **D)** S phase (determined by BrdU incorporation and ELISA) in UL141 (or control) expressing HUVECs in the presence/absence of increasing amounts of the topoisomerase inhibitor, camptothecin (CPT). Data represents 3 independent experiments performed in triplicate. **E)** Immunofluorescence microscopy of HUVECs expressing UL141 (or control) was used to assess p53 expression and cell surface expression of PVR, with/without Nutlin-3 treatment for 18 h; quantitation of nuclear p53 is shown in panel F.**F)** Quantitation of nuclear p53 from panel E. Data was obtained from 3 independent experiments performed in duplicate **G)** UL141 (or vector only) transduced HUVECs were treated with combinations of siRNAs targeting PVR, TP53 and/or MDM2 for 72 h before assessment of protein expression by western blotting.

The absence of p53 induction in UL141-expresing HUVECs suggested that intracellular PVR continues to deliver the signals that maintain p53 turnover. To test this, we depleted PVR (using siRNA) from both control and UL141-expressing HUVECs (Fig 4*G)*; PVR depletion from the UL141-expressing HUVECs induced p53 and its downstream targets. Accordingly, induction of p21 and MDM2 in PVR depleted, UL141-expressing HUVECs was p53 dependent (Fig. 4*G*). Thus, in HUVECs, PVR is still capable of limiting p53 induction when retained within the ER by UL141, thereby allowing HCMV to evade both immune surveillance and to prevent activation of the p53-PVR axis. However, the PVR-p53 signalling axis signalling works most efficiently from the cell surface and UL141-expressing HUVECs exhibited enhanced p53-dependent, activation of caspase 3 cleavage (Fig. 4*G*). Whilst UL141 expression in HUVECs did not alter sensitivity to either nutlin-3 or the topoisomerase I inhibitor camptothecin (Fig. 4*D-F*), it did sensitise cells to the action of doxorubicin, a topoisomerase II inhibitor; UL141-expressing HUVECs showed significantly reduced viability in the presence of doxorubicin and this was associated with increased accumulation of p53 in the nucleus (Supplementary Fig. *4E-G*). This suggests that UL141 expression (and PVR retention in the ER) imparts a low level of stress which in itself is insufficient to induce p53. However, this stress sensitises HUVECs to other p53-inducing agents and demonstrates that the PVR-p53 axis acts collectively with other signals to regulate p53 activity.

## Discussion

Various types of stress induce signals that result in the stabilisation of p53 and the induction of its downstream transcriptional programme. This system has evolved to limit cell proliferation under sub-optimal conditions, a situation that is likely to result in the accumulation of deleterious mutations, with implications for developmental pathways and cancer. Amongst the sources of cell stress is the prototypical p53 inducer, genotoxic stress, along with nutrient deprivation, redox status and hypoxia (1–3). We have identified an additional trigger for p53 induction in the endothelium, the loss of cell surface PVR.

The PVR molecule (CD155/NECL5) is multifunctional, with roles in cell adhesion, migration and growth/contact inhibition, as well as immune regulation (19–23, 28, 31–33, 35). For EC, PVR is important for the adhesion and transendothelial migration (TEM) of leucocytes during inflammation (19, 35). Mice lacking *Cd155* are viable and fertile, but demonstrate impaired angiogenesis following hindlimb ischemia; *Cd155*^-/-^ ECs proliferate poorly *ex vivo* and RNAi of CD155 in HUVECs resulted in a reduction in cells in S phase and impaired angiogenic activity (47). Furthermore, the same study identified a physical interaction between transfected, epitope tagged CD155 and tagged VEGFR2. We show that the impairment of EC proliferation and angiogenic functions induced upon PVR loss is p53 dependent, linking PVR to the established critical role of p53 in angiogenesis (39). Use of UL141 and cell surface biotinylation showed that UL141 removes endogenous PVR from the HUVEC cell surface without loss of surface VEGFR2, indicating that PVR is not simply regulating VEGFR2 expression. Our data show that, under steady-state conditions, PVR actively maintains p53 turnover and that loss of cell surface PVR expression results in reduced p53 ubiquitination, p53 stabilisation and the induction of downstream pathways. Previously, nutlin-3 treatment has been shown to induce p53 via disruption of MDM2 mediated ubiquitination (48). In ECs, this culminated in angiogenic defects, highlighting the key role that p53 plays in controlling EC growth and differentiation (39). However, whilst nutlin-3 disrupts MDM2-p53 interactions, we found that PVR loss reduced p53 ubiquitination without impeding MDM2 binding, suggesting a potential role for deubiquitinase enzymes (DUBs) in coordinating the PVR-p53 axis (49). Loss of PVR may activate DUBs implicated in p53 control, such as USP3, 7, 10, and 13, to deubiquitinate p53 and promote its nuclear accumulation (49). Alternatively, disruption of the ubiquitin ligase conjugating systems controlling p53 ubiquitination, at either the E2, or E3 level, would also prevent ubiquitin addition and reduce proteasomal turnover of p53 (50). Membrane-associated RING-CH-type (MARCH) finger 7 ubiquitin ligase catalyses polyubiquitination of MDM2 (51). This limits MDM2 autoubiquitination, thereby stabilising MDM2 levels and promoting p53 ubiquitination and turnover. Thus, PVR loss could suppress MARCH7 ubiquitin ligase activity, resulting in increased MDM2 turnover and limiting the proteasomal degradation of p53. In addition, the importance of post-translational modifications in the control of MDM2 and p53 stability and function is well-established (52). Acetylation of MDM2 also antagonizes autoubiquitination, promoting deubiquitination and increasing its ubiquitin ligase activity against p53 (53, 54). Deacetylation of MDM2 by sirtuin 1 (SIRT1) promotes MDM2 degradation and stabilises p53 protein levels (53). Thus, PVR may alternatively dampen SIRT1 deacetylase activity under normal conditions, but upon down-regulation, this suppression is released to ultimately stabilise p53 protein levels. The precise signals linking PVR expression to the control of p53 ubiquitination is currently under investigation.

The PVR molecule is an important regulator of immune function. The induction of PVR expression in malignant or infected cells provides a signal to trigger cellular immunity via the activating receptor, DNAM-1. Many viruses encode molecules that modulate the expression of DNAM-1 ligands, aiding evasion of NK cells and, for HCMV, UL141 blocks cell surface expression of the two known DNAM-1 ligands, PVR and nectin-2/CD112 (28, 30). Our use of HCMV UL141 was intended primarily as a tool to prevent PVR expression at the cell surface.

However, our data showing that UL141-mediated retention of PVR maintains p53 turnover provides insight into the action of UL141 and suggests a physiological function for the PVR-p53 axis. The UL141 molecule retains both PVR and nectin-2 in the ER, but whilst PVR is held intact in this compartment, nectin-2 is degraded by the action of HCMV US2 (30, 45). The different fates of PVR and nectin-2 in HCMV infected cells was puzzling. However, our results show that PVR expression must be maintained to prevent p53 stabilisation, cell cycle arrest and apoptosis. We suggest that UL141 has evolved to retain intact PVR in the ER for two reasons, to prevent recognition of infected cells by NK cells and to retain p53 turnover. Thus, PVR contributes two surveillance functions; in a cell extrinsic pathway, PVR expression at the cell surface acts as a signal for immune recognition via DNAM-1 (20), eliminating infected or defective cells. In the cell intrinsic pathway identified here, PVR downregulation in various EC types induces p53 stabilisation and cell cycle arrest. The ability to protect against PVR loss via p53 stabilisation provides a mechanism to counteract pathogen immune evasion strategies targeting PVR, albeit one that HCMV has subverted. The expression of UL141 in EC will presumably allow HCMV to evade both the cell extrinsic and cell intrinsic sensor functions of PVR, contributing to HCMV associated vascular disease and favouring EC as a site of viral persistence (55–58). Furthermore, PVR is required for endothelial proliferation, migration, and the formation of capillary networks. Triggering of a p53-mediated arrest program following PVR loss will minimise the production of immature blood vessels lacking this multifunctional molecule; such structures would be expected to have altered adhesion and permeability to both extravasating and intravasating cells. Such activity might be restricted to vascular repair, since the PVR molecule is not required for development in mice, but is important in maintaining physiological function in adults after injury (47).

The PVR molecule activates NK cells and T cells via DNAM-1 and *Dnam1-/-* mice show a reduced ability to control tumours (59). However, PVR has two additional receptors, TIGIT and CD96 which, unlike DNAM-1, deliver inhibitory signals (23). This interaction of CD155 with TIGIT and/or CD96 provides immune checkpoint activity that limits anti-tumour responses. Furthermore, *Cd155*^-/-^ mice exhibit enhanced graft versus host disease and expression of CD155 on human ECs inhibits T cell activity (60, 61). The PVR-p53 axis might ensure that EC are capable of delivering these inhibitory signals under inflammatory conditions, minimising damage to the healthy endothelium. Whether the PVR-p53 axis operates in other cell types is currently unknown. The high levels of PVR expression seen in many tumour types is predicted to favour tumour growth, the loss of contact inhibition and migratory activity (31, 33, 47, 62, 63). The ability of highly expressed PVR to promote p53 turnover will contribute to malignancy, although this is only likely to be of importance in tumours with an intact p53 pathway.

In summary, we have identified loss of EC surface PVR as a trigger for p53 stabilisation and concomitant cell cycle arrest. Expression of PVR provides a rheostat for EC health, with low expression activating the p53 pathway and high expression being a signal for immune recognition. This pathway is subverted by the HCMV UL141 molecule, allowing this virus to escape the cell extrinsic immune surveillance functions of PVR and the cell intrinsic PVR-p53 axis identified here. Importantly, this work identifies a previously unknown role for p53. From its originally identified role in protecting genome integrity (1), p53 has emerged as a critical hub for regulating the responses to diverse types of cellular stress, including DNA damage, hypoxia, oxidative damage and various metabolic demands (2, 3). Our data identify p53 as critical to monitoring cell surface composition, a role that bridges the functions of p53 in cellular stress and its involvement in healthy proliferation and differentiation pathways (3, 15).

## Materials and Methods

### Antibodies and associated reagents

The following antibodies were used for flow cytometry, immunofluorescence microscopy, immunoprecipitation and immunoblotting. Anti-PVR (SKII.4), anti-CD146-APC (SHM-57 or P1H12), anti-CD31-AF488 (WM-59), anti-CD99 (HCD99) were from BioLegend. Anti-cyclin A (BF683), anti-cyclin B (GNS-11), anti-CDK1 (clone 1), anti-CDK2 (clone 55), anti-CDH5-Biotin (clone 75), anti-CD45-FITC (clone 69), were from BD Biosciences. Rabbit Anti-PVR (D3G7H), Anti-VEGFR2 (D5B1), Anti-caspase 3 (8G10), anti-cleaved caspase 3 (5A1E), anti-PARP (#9542), anti-Bax (#2772), anti-HAUSP (#3277), anti-CDK1 (POH1), anti-phospho-ERK1/2 ((#9106; E10), anti-V5 tag (#13202; D3H8Q), anti-phospho-S6 (5364; D68F9), anti-phospho-AKT (Ser473; #4060; D9E), and anti-CDK2 (78B2) were from Cell Signalling Technologies. Anti-p53 (DO-1, pAB1801, FL393), anti-p21 (C20; C-19; SC-397-G), anti-MDM2 (SPM14), anti-VE-cadherin (BV9), anti-PCNA (PC10), anti-pRB (H-2), and anti-cyclin A (H-432) were from Santa Cruz Biotechnology. Anti-Nectin-2 (AF2229), anti-ubiquitin (FK2), and anti-VEGFR2 (AF357) were from R&D Technologies. Anti-GFP (G1544) and Anti-Actin (A544a; AC-15) were from Sigma. Anti-HSP90 (4C10), anti-GAPDH (2D9) and a rabbitt polyclonal anti-CEACAM1 (TA350817) antibody were from Origene. Anti-Rabbit-HRP and anti-mouse-HRP were from Cell Signalling Technologies. All AlexaFluor-conjugated secondary antibodies were from Life Technologies (ThermoFisher). Streptavidin-HRP was from DAKO.

### Cell culture

Primary endothelial cells were cultured in complete Endothelial Growth Media (EGM or EGM MV2; Promocell) and were maintained on either gelatin (HUVEC, HBMEC), fibronectin (HDLEC), or collagen (BOEC, hCMEC/D3) coated dishes. HUVECs, HBMEC and HDLEC (neonatal) cells were obtained from Promocell and used up until passage 6. The hCMEC/D3 cell line was obtained from VHBio Ltd (CLU512 from Cedarlane) at passage 26 and used until passage 35. Blood outgrowth endothelial cells were isolated from whole blood as described (64). Briefly, whole blood was overlaid onto Lymphoprep and spun for 15 minutes at 450 x g without braking before removal of the buffy coat. Cells were diluted in PBS before centrifugation at 600 x g for 7 minutes and resuspension in ECGM MV2 supplemented with Lipid Supplement, Insulin and hydrocortisone. Cells were allowed to adhere to type I collagen-coated dishes overnight prior to washing with PBS containing 10% FCS and replacement with fresh modified MV2 media. Media was replaced every 3 days for 3 weeks before the appearance of colonies and subsequent BOEC expansion. Cells were used between passage 5 and 13. Once expanded, all endothelial cells (HUVEC, hCMEC/D3, HDLEC, HDBEC, and BOECs) were maintained in either ECGM or ECGM MV2 (Promocell) and endothelial phenotype regularly assessed by flow cytometry using anti-CD31-AF488 and anti-VE-cadherin-Biotin antibodies. Primary and hTERT immortalised human dermal foreskin fibroblasts (HDFs), PC-3, U251 and SW620 cells were maintained in DMEM supplemented with 10% FCS. MDA-MB-231, HL60 cells and primary PBMCs were maintained in RPMI supplemented with 10% FCS.

### Primers

*TP53; For: CCTCAGCATCTTATCCGAGTGG; Rev: TGGATGGTGGTACAGTCAGAGC MDM2; For: TGTTTGGCGTGCCAAGCTTCTC; Rev: CACAGATGTACCTGAGTCCGATG CDKN1A; For: AGGTGGACCTGGAGACTCTCAG; Rev: TCCTCTTGGAGAAGATCAGCCG Bax; For: TCAGGATGCGTCCACCAAGAAG; Rev: TGTGTCCACGGCGGCAATCATC CDK1; For: GGAAACCAGGAAGCCTAGCATC; Rev: GGATGATTCAGTGCCATTTTGCC CDK2; For: ATGGATGCCTCTGCTCTCACTG; Rev: CCCGATGAGAATGGCAGAAAGC CCNA2; For: CTCTACACAGTCACGGGACAAAG; Rev: CTGTGGTGCTTTGAGGTAGGTC CCNB1; For: GACCTGTGTCAGGCTTTCTCTG; Rev: GTATTTTGGTCTGACTGCTTGC CCND1; For: TCTACACCGACAACTCCATCCG; Rev: TCTGGCATTTTGGAGAGGAAGTG PVR; For: CACTGTCACCAGCCTCTGGATA; Rev: TCATAGCCAGAGATGGATACCTC MCAM; For: ATCGCTGCTGAGTGAACCACAG; Rev: CTACTCTCTGCCTCACAGGTCA*

### siRNA transfection

Cells were reverse transfected with SMARTpool or individual siRNA duplexes (Dharmacon or Ambion) using either Lipofectamine RNAiMAX (Invitrogen) or Dharmafect I (Dharmacon) according to manufacturer’s instructions and as outlined previously (37). Briefly 24 pmol siRNA was used together with 4 µl of liposomal transfection reagent in 400 µl OptiMEM (Invitrogen) per 3×10^5^ cells and scaled accordingly. Cell and transfection mixtures were incubated for 4h in OptiMEM before replacement with complete growth media. Cells were used in downstream assays following 24-72 h culture as indicated. ON-TARGETplus Human PVR (5817) siRNA – SMARTpool (5’-GGAUCGGGAUUUAUUUCUA-3’, 5’-CCAAACGGCUGGAAUUCGU-3’, 5’-GGGCAUGUCUCCUAUUCAG-3’, 5’-GCAAGAAUGUGACCUGCAA-3’); Scrambled (ON-TARGETplus Non-targeting control pool, Dharmacon; D-001810); p53 (Ambion; s637); and MDM2 (Ambion; s8630).

### Viral transduction

Lentiviral supernatants were produced in HEK293Lx cells following transfection with 3^rd^ generation packaging plasmids and either pLKOshp53 (Addgene #19119), pLKOshScr (Addgene #1864), pshPVR (ABM #i018439) for shRNA or pHAGE-Scr or pHAGE-UL141 for heterogenous expression. Transfectants were selected using the Puromycin selection marker before use as a pooled population. Experiments were repeated using cells from independent transductions. For HCMV infections of primary human dermal fibroblasts, the Merlin virus strain harboring modified GFP-E6 protein was produced in human dermal fibroblasts, titred and used at MOI of 1 or 10 to infect cells for 72-96h before assaying protein expression by flow cytometry and Western blotting.

### Proliferation, toxicity and migration assays

For BrdU incorporation, siRNA transfected cells were counted and reseeded in 96 well plates for 24 h prior to incubation with 10 µM BrdU for 4 h and fixation in 70% ethanol. Nuclei were denatured in 2N HCl/1% TX100 before successive incubations with anti-BrdU-Biotin antibodies (BioLegend) and streptavidin-HRP (DAKO) prior to development with TMB. Cell viability was determined by MTT assay according to manufacturer’s instructions (Sigma). Scratch wound assays were performed using a 96 well plate scratch tool and monitored using an Incucyte imaging system. Transwell migration assays were undertaken using 5 µm, 13mm diameter filters (BD) for 4-18h before fixation of filters and staining for migrated cells using phalloidin, Hoecsht 33342, or CD31/VE-cadherin staining. For analysis of PBMC transendothelial migration, leukocytes were pre-labelled with CellTracker Green (Life Technologies) prior to addition to Transwell filters containing confluent endothelial cells. Migrated cells were collected from the bottom well, stained with anti-CD3 (BioLegend) and anti-CD45 (BD) antibodies and counted using an Attune flow cytometer. Endothelial bound PBMCs were also assayed by staining and imaging the upper and bottom Transwell filters.

### Western blotting, biotinylation and immunoprecipitation

Protein expression was assessed using SDS–polyacrylamide gel electrophoresis (PAGE). Cells were washed three times with PBS and lysed in 2% SDS lysis buffer as outlined previously. Protein concentration was determined by bicinchoninic acid assay (ThermoFisher) and 15-25 µg total protein separated on a 5-20% polyacrylamide gradient gels prior to Western blotting on Nitrocellulose membranes. Horseradish peroxidase (HRP)-conjugated secondary antibodies were used for detection by enhanced chemiluminescence (ECL) (BioRad). Chemiluminescence was detected either using ECL Hyperfilm (GEHealthcare) or using a ChemiDoc imager (BioRad). Protein quantitation was performed using either ImageLab (BioRad; Ver 5.2.1) or ImageStudioLite (Ver 5.2.5; LI-COR). For cell surface biotinylation, EC were washed three-times with ice-cold PBS and incubated with 0.3 mg/ml EZ-link Sulfo-NHS-LC-Biotin (ThermoFisher) in PBS containing 2 mM MgCl2 and 2 mM CaCl2 on ice for 30 minutes with gentle agitation. Biotinylated cells were then quenched in TBS (20 mM Tris pH 7.6, 137 mM NaCl) followed by 2 washes in PBS prior to lysis in RIPA buffer (1% TX100, 50 mM Tris/HCl (pH 7.6), 150 mM NaCl, 1 mM EDTA, 1 mM EGTA, 0.25% sodium deoxycholate, 0.05% SDS with protease and phosphatase inhibitors). Cleared lysates were subjected to BCA assay (ThermoFisher) before incubation of equalised protein concentrations with 30 μl neutravidin-agarose beads (Pierce) for 2 hours at 4°C. Washed beads were analysed by Western blotting for bound proteins. For immunoprecipitations, 500 µg of precleared total protein lysates of indicated cells were generated in modified 0.5x RIPA buffer (see above) and incubated with 25 µl Protein A/G agarose and 2 µg antibody. Beads were washed in 4 times in lysis buffer before analysis of bound proteins by Western blotting

### Immunofluorescence microscopy

5×10^4^ endothelial cells were seeded to 0.2% gelatin-coated 13mm diameter coverslips (VWR) or 1×10^4^ cells seeded to gelatin-coated 96 well ViewPlates (Perkin Elmer) and grown to confluency. Cells were washed once in PBS prior to fixation in 4% paraformaldehyde and permeabilisation with 0.2% TritonX100 in PBS. Primary antibody staining was performed overnight in either PBLEC (PBS with 2mM CaCl2, 2mm MgCl2, 0.1 mM MnCl, 1% Tween 20, 0.2% BSA, 0.2% FCS) or 2% horse serum in PBS followed by conjugated secondary antibodies for 1 h before mounting onto slides with Fluoromount G. For PVR staining, cells were labelled with anti-PVR (SKII.4) antibody in 0.5% BSA in PBS (with Ca^2+^ and Mg^2+^) for 20 min prior fixation in 4% paraformaldehyde and processing as above. Subsequent imaging was carried out using an Essen Incucyte Zoom live cell imager (Essen Bioscience), Operetta HTS imager (Perkin Elmer), EVOS miscroscope system (Thermo Fisher Scientific), or Nikon A1R confocal microscope (Nikon). Image analysis was performed using ImageJ (https://imagej.nih.gov/ij/) or Columbus (Perkin Elmer) software.

### Flow cytometry

siRNA pre-treated endothelial cells were detached using Accutase (LifeTechnologies, ThermoFisher) and resuspended in DMEM containing 10% FCS. Cells were subsequently spun down and washed 2x in PBS/0.2% BSA before live cell staining. Cells were incubated with indicated fluorescently-conjugated antibodies on ice for 20-30 minutes in staining buffer (2% BSA or 1% FCS, 0.05% NaN3 in PBS). Analysis was performed on either an LSRII 3 or 4-laser flow cytometer (Becton Dickinson) or Attune Acoustic Flow Cytometer (Roche). Post analysis was performed using either FACSDiva (BD), or FlowJo version X (BD).

### Collagen overlay assay

For short term assessment of endothelial tube formation, collagen overlay of HDLECs was performed as described previously (37). Briefly, siRNA reverse transfected cells were grown to confluency before overlaying with neutralised type I collagen (1 mg/ml; Rat tail; Millipore). Collagen was allowed to set for 45 minutes before overlaying with complete growth media. Following 18-24 hours incubation at 37°C, 5% CO2, cells were fixed in 4% paraformaldehyde, stained with phalloidin (LifeTechnologies; ThermoFisher) and Hoechst 33342 before imaging using an Essen Incucyte Zoom live cell imager (Essen Bioscience) or Operetta HTS Imager. Tubules were quantified using either ImageJ or Essen Incucyte Zoom software.

### Tubulogenesis assays

ECs were reverse-transfected with indicated SMARTpool siRNA or scrambled control siRNA (ON-TARGETplus Non-targeting control pool, Dharmacon). Tubulogenesis assays were performed as described (38). HDFs were seeded at a density of 2×10^4^/well. The following day, 2.5×10^3^ siRNA treated endothelial cells were seeded to confluent HDF feeder layers in ECBM. Media was then replaced every 2 days with complete growth media supplemented with 10 ng/ml VEGFA. Co-cultures were fixed at 7-10 days in either 4% paraformaldehyde for 10 minutes or ice-cold 70% ethanol for 30 minutes prior to staining with anti-CD31-AF488-conjugated antibody (BioLegend) and anti-VWF (DAKO) followed by secondary donkey anti-mouse AF488 and donkey anti-rabbit AF555 antibodies. Images were obtained using an Essen Incucyte Zoom live cell imager (Essen Bioscience) 4x objective and analysis performed using ImageJ (Angiogenesis Analyzer plug-in), AngioTool64 (http://angiotool.nci.nih.gov), or Essen Incucyte program.

### Peripheral Blood Mononuclear Cell (PBMC) isolation and adhesion assay

Waste apheresis samples (from healthy donors) were purchased from NHS Blood and Transplant. Peripheral blood mononuclear cells were isolated using the density gradient medium Lymphoprep (StemCell Technologies). Primary isolated PBMCs were Cell Tracker Green (CTG) (Thermo Fisher) labelled at 4×10^6^ cells/mL for 30 minutes at 37°C 5% CO2. Cells were then pelleted and washed twice in RPMI before being resuspended at 1×10^6^/mL. CTO-labelled PBMCs were seeded to confluent HUVEC monolayers at a ratio of 5:1 and left to adhere for 1 hour at 37°C. Cells were washed 1x in PBS and fixed in 4% PFA before further gentle washing with PBS. Images were obtained using Operetta high content imaging system (Perkin Elmer) and counted with Columbus software.

### xCelligence RTCA

Real-time monitoring of growth curves and cellular migration were measured using an electrical impedence based assay using the xCELLigence RTCA DP instrument (ACEA Biosciences, Inc.). E-plates/CIM plates were coated prior to plating endothelial cells, with human fibronectin (Sigma) 10 μg/ml for 2h at room temperature under a laminar flow hood. Wells were then rinsed 2x with PBS and air dried before seeding of cells. For assessment of growth rates, (2×10^3^-5×10^3^) siRNA reverse transfected endothelial cells (24h or 48h post transfection) were seeded to fibronectin (Sigma)-coated E-plates (growth) or CIM plates (migration) (ACEA Biosciences, Inc.) and analysed using xCELLigence RTCA DP instrument. Impedance was measured every 15 minutes for 48-96 hours. For real-time monitoring of cellular migration, CIM-plate 16 inserts (ACEA Biosciences Inc.) were used where endothelial cells were plated in serum-free media to the upper chamber and the lower chamber loaded with 10% FCS-containing complete growth media as a chemoattractant. Impedance was measured every 15 minutes for 72 hours.

### Gene expression analysis

Total RNA was isolated from various ECs using RNeasy Plus Kit according to manufacturer’s instructions (Qiagen). 200-500 ng RNA was reverse transcribed using High-Capacity RNA-to-cDNA Kit (ThermoFisherScientific) prior to gene expression analysis by SYBRGreen-based QPCR (Amersham 7900HT or Q5 systems), using GAPDH as an internal control. Data was analysed using deltadeltaCt calculations for the test and GAPDH genes.

## Acknowledgments

This work was supported by the University of Leeds. We are grateful to Peter Tomasec, Luis Nobre and Michael Weekes for reagents and to those colleagues who provided comments on the work.

## Author contribution statement

AFO, PJF and GPC conceived the study. AFO and AJM performed the experiments. All authors analysed and discussed the data and all authors contributed to the writing and editing of the manuscript. PFJ and GPC made an equal contribution to the work.

## Declaration

The authors declare no competing interests.

## References

1. D. P. Lane, Cancer. p53, guardian of the genome. Nature 358, 15–16 (1992).

2. F. Kruiswijk, C. F. Labuschagne, K. H. Vousden, p53 in survival, death and metabolic health: a lifeguard with a licence to kill. Nat Rev Mol Cell Biol 16, 393–405 (2015).

3. E. R. Kastenhuber, S. W. Lowe, Putting p53 in Context. Cell 170, 1062–1078 (2017).

4. A. Efeyan, M. Serrano, p53: Guardian of the genome and policeman of the oncogenes. Cell Cycle 6, 1006–1010 (2007).

5. S. Y. Shieh, J. Ahn, K. Tamai, Y. Taya, C. Prives, The human homologs of checkpoint kinases Chk1 and Cds1 (Chk2) phosphorylate p53 at multiple DNA damage-inducible sites. Genes Dev. 14, 289–300 (2000).

6. M. H. Kubbutat, S. N. Jones, K. H. Vousden, Regulation of p53 stability by Mdm2. Nature 387, 299–303 (1997).

7. Y. Haupt, R. Maya, A. Kazaz, M. Oren, Mdm2 promotes the rapid degradation of p53. Nature 387, 296–299 (1997).

8. T. Riley, E. Sontag, P. Chen, A. Levine, Transcriptional control of human p53-regulated genes. Nat Rev Mol Cell Biol 9, 402–412 (2008).

9. A. Hafner, M. L. Bulyk, A. Jambhekar, G. Lahav, The multiple mechanisms that regulate p53 activity and cell fate. Nat. Rev. Mol. Cell Biol. 20, 199–210 (2019).

10. W. S. el-Deiry, et al., WAF1, a potential mediator of p53 tumor suppression. Cell 75, 817–825 (1993).

11. J. Wade Harper, G. R. Adami, N. Wei, K. Keyomarsi, S. J. Elledge, The p21 Cdk-interacting protein Cip1 is a potent inhibitor of G1 cyclin-dependent kinases. Cell 75, 805–816 (1993).

12. Y. Barak, T. Juven, R. Haffner, M. Oren, Mdm2 Expression Is Induced By Wild Type P53 Activity. EMBO J. 12, 461–468 (1993).

13. T. Miyashita, J. C. Reed, Tumor suppressor p53 is a direct transcriptional activator of the human bax gene. Cell 80, 293–299 (1995).

14. J. Reyes, et al., Fluctuations in p53 Signaling Allow Escape from Cell-Cycle Arrest. Mol. Cell 71, 581-591.e5 (2018).

15. N. Rivlin, G. Koifman, V. Rotter, p53 orchestrates between normal differentiation and cancer. Semin Cancer Biol 32, 10–17 (2015).

16. L. A. Donehower, et al., Integrated Analysis of TP53 Gene and Pathway Alterations in The Cancer Genome Atlas. Cell Rep. 28, 1370–1384.e5 (2019).

17. M. L. Tornesello, C. Annunziata, A. L. Tornesello, L. Buonaguro, F. M. Buonaguro, Human oncoviruses and p53 tumor suppressor pathway deregulation at the origin of human cancers. Cancers (Basel). 10 (2018).

18. C. L. Mendelsohn, E. Wimmer, V. R. Racaniello, Cellular receptor for poliovirus: molecular cloning, nucleotide sequence, and expression of a new member of the immunoglobulin superfamily. Cell 56, 855–865 (1989).

19. N. Reymond, et al., DNAM-1 and PVR regulate monocyte migration through endothelial junctions. J Exp Med 199, 1331–1341 (2004).

20. C. Bottino, et al., Identification of PVR (CD155) and Nectin-2 (CD112) as cell surface ligands for the human DNAM-1 (CD226) activating molecule. J Exp Med 198, 557–567 (2003).

21. S. Tahara-Hanaoka, et al., Functional characterization of DNAM-1 (CD226) interaction with its ligands PVR (CD155) and nectin-2 (PRR-2/CD112). Int. Immunol. 16, 533–538 (2004).

22. C. J. Chan, et al., DNAM-1/CD155 Interactions Promote Cytokine and NK Cell-Mediated Suppression of Poorly Immunogenic Melanoma Metastases. J. Immunol. 184, 902–911 (2010).

23. J. S. O’Donnell, J. Madore, X. Y. Li, M. J. Smyth, Tumor intrinsic and extrinsic immune functions of CD155. Semin. Cancer Biol. (2020) https://doi.org/10.1016/j.semcancer.2019.11.013.

24. T. Hirota, K. Irie, R. Okamoto, W. Ikeda, Y. Takai, Transcriptional activation of the mouse Necl-5/Tage4/PVR/CD155 gene by fibroblast growth factor or oncogenic Ras through the Raf-MEK-ERK-AP-1 pathway. Oncogene 24, 2229–2235 (2005).

25. A. Soriani, et al., ATM-ATR-dependent up-regulation of DNAM-1 and NKG2D ligands on multiple myeloma cells by therapeutic agents results in enhanced NK-cell susceptibility and is associated with a senescent phenotype. Blood 113, 3503–3511 (2009).

26. N. Kamran, et al., Toll-Like Receptor Ligands Induce Expression of the Costimulatory Molecule CD155 on Antigen-Presenting Cells. PLoS One 8, e54406 (2013).

27. D. Pende, et al., Expression of the DNAM-1 ligands, Nectin-2 (CD112) and poliovirus receptor (CD155), on dendritic cells: Relevance for natural killer-dendritic cell interaction. Blood 107, 2030–2036 (2006).

28. P. Tomasec, et al., Downregulation of natural killer cell-activating ligand CD155 by human cytomegalovirus UL141. Nat Immunol 6, 181–188 (2005).

29. G. Matusali, M. Potestà, A. Santoni, C. Cerboni, M. Doria, The Human Immunodeficiency Virus Type 1 Nef and Vpu Proteins Downregulate the Natural Killer Cell-Activating Ligand PVR. J. Virol. 86, 4496–4504 (2012).

30. V. Prod’homme, et al., Human cytomegalovirus UL141 promotes efficient downregulation of the natural killer cell activating ligand CD112. J Gen Virol 91, 2034–2039 (2010).

31. T. Fujito, et al., Inhibition of cell movement and proliferation by cell-cell contact-induced interaction of Necl-5 with nectin-3. J Cell Biol 171, 165–173 (2005).

32. Y. Takai, J. Miyoshi, W. Ikeda, H. Ogita, Nectins and nectin-like molecules: roles in contact inhibition of cell movement and proliferation. Nat Rev Mol Cell Biol 9, 603–615 (2008).

33. W. Ikeda, et al., Nectin-like molecule-5/Tage4 enhances cell migration in an integrin-dependent, Nectin-3-independent manner. J Biol Chem 279, 18015–18025 (2004).

34. B. B. Weksler, et al., Blood-brain barrier-specific properties of a human adult brain endothelial cell line. FASEB J. 19, 1872–1874 (2005).

35. D. P. Sullivan, M. A. Seidman, W. A. Muller, Poliovirus receptor (CD155) regulates a step in transendothelial migration between PECAM and CD99. Am J Pathol 182, 1031–1042 (2013).

36. S. P. Herbert, D. Y. R. Stainier, Molecular control of endothelial cell behaviour during blood vessel morphogenesis. Nat. Rev. Mol. Cell Biol. 12, 551–564 (2011).

37. S. P. Williams, et al., Genome-wide functional analysis reveals central signaling regulators of lymphatic endothelial cell migration and remodeling. Sci. Signal. 10, eaal2987. (2017).

38. E. T. Bishop, et al., An in vitro model of angiogenesis: basic features. Angiogenesis 3, 335–344 (1999).

39. P. Secchiero, et al., Antiangiogenic activity of the MDM2 antagonist nutlin-3. Circ Res 100, 61–69 (2007).

40. X. Wu, J. H. Bayle, D. Olson, A. J. Levine, The p53-mdm-2 autoregulatory feedback loop. Genes Dev 7, 1126–1132 (1993).

41. M. Li, et al., Deubiquitination of p53 by HAUSP is an important pathway for p53 stabilization. Nature 416, 648–653 (2002).

42. M. Fischer, Census and evaluation of p53 target genes. Oncogene 36, 3943–3956 (2017).

43. S. E. Jackson, G. M. Mason, M. R. Wills, Human cytomegalovirus immunity and immune evasion. Virus Res. 157, 151–160 (2011).

44. M. Patel, et al., HCMV-encoded NK modulators: Lessons from in vitro and in vivo genetic variation. Front. Immunol. 9 (2018).

45. J. L. Hsu, et al., Plasma membrane profiling defines an expanded class of cell surface proteins selectively targeted for degradation by HCMV US2 in cooperation with UL141. PLoS Pathog 11, e1004811 (2015).

46. P. Muganda, O. Mendoza, J. Hernandez, Q. Qian, Human cytomegalovirus elevates levels of the cellular protein p53 in infected fibroblasts. J Virol 68, 8028–8034 (1994).

47. M. Kinugasa, et al., Necl-5/poliovirus receptor interacts with VEGFR2 and regulates VEGF-induced angiogenesis. Circ. Res. 110, 716–726 (2012).

48. L. T. Vassilev, et al., In Vivo Activation of the p53 Pathway by Small-Molecule Antagonists of MDM2. Science (80-.). 303, 844–848 (2004).

49. S. K. Kwon, M. Saindane, K. H. Baek, p53 stability is regulated by diverse deubiquitinating enzymes. Biochim. Biophys. Acta -Rev. Cancer 1868, 404–411 (2017).

50. I. Gupta, K. Singh, N. K. Varshney, S. Khan, Delineating crosstalk mechanisms of the ubiquitin proteasome system that regulate apoptosis. Front. Cell Dev. Biol. (2018) https://doi.org/10.3389/fcell.2018.00011.

51. K. Zhao, et al., Regulation of the Mdm2–p53 pathway by the ubiquitin E3 ligase MARCH 7. EMBO Rep. (2018) https://doi.org/10.15252/embr.201744465.

52. M. F. Lavin, N. Gueven, The complexity of p53 stabilization and activation. Cell Death Differ. (2006) https://doi.org/10.1038/sj.cdd.4401925.

53. N. T. Nihira, et al., Acetylation-dependent regulation of MDM2 E3 ligase activity dictates its oncogenic function. Sci. Signal. (2017) https://doi.org/10.1126/scisignal.aai8026.

54. N. Patel, et al., HDAC2 Regulates Site-Specific Acetylation of MDM2 and Its Ubiquitination Signaling in Tumor Suppression. iScience 13, 43–54 (2019).

55. Y. H. Shen, et al., Human cytomegalovirus causes endothelial injury through the ataxia telangiectasia mutant and p53 DNA damage signaling pathways. Circ Res 94, 1310–1317 (2004).

56. B. Adler, C. Sinzger, Endothelial cells in human cytomegalovirus infection: one host cell out of many or a crucial target for virus spread? Thromb Haemost 102, 1057–1063 (2009).

57. F. Bughio, M. Umashankar, J. Wilson, F. Goodrum, Human Cytomegalovirus UL135 and UL136 Genes Are Required for Postentry Tropism in Endothelial Cells. J Virol 89, 6536–6550 (2015).

58. M. A. Jarvis, J. A. Nelson, Human cytomegalovirus tropism for endothelial cells: not all endothelial cells are created equal. J Virol 81, 2095–2101 (2007).

59. A. Iguchi-Manaka, et al., Accelerated tumor growth in mice deficient in DNAM-1 receptor. J. Exp. Med. 205, 2959–2964 (2008).

60. S. Seth, et al., Absence of CD155 aggravates acute graft-versus-host disease. Proc. Natl. Acad. Sci. U. S. A. 108 (2011).

61. N. K. Escalante, A. Von Rossum, M. Lee, J. C. Choy, CD155 on human vascular endothelial cells attenuates the acquisition of effector functions in CD8 T cells. Arterioscler. Thromb. Vasc. Biol. 31, 1177–1184 (2011).

62. S. Kakunaga, et al., Enhancement of serum- and platelet-derived growth factor-induced cell proliferation by Necl-5/Tage4/poliovirus receptor/CD155 through the Ras-Raf-MEK-ERK signaling. J. Biol. Chem. 279, 36419–36425 (2004).

63. K. E. Sloan, et al., CD155/PVR plays a key role in cell motility during tumor cell invasion and migration. BMC Cancer (2004) https://doi.org/10.1186/1471-2407-4-73.

64. J. Martin-Ramirez, M. Hofman, M. van den Biggelaar, R. P. Hebbel, J. Voorberg, Establishment of outgrowth endothelial cells from peripheral blood. Nat Protoc 7, 1709–1715 (2012).

