## Supplementary Figures and Legends for "Negative regulation of p53 by the poliovirus receptor PVR is a target of a human cytomegalovirus immune evasion molecule"

#### Supplementary Figure 1

**A)** Confluent (Conf) and sub-confluent (Sub-conf) endothelial cells, leukocytes, and cell lines (as used in Figure 1) were assessed for PVR and ICAM-1 expression by flow cytometry after growth for 18 h in either serum-free (SF) or full growth media (FM). **B)** Quantitation of PVR and **C)** ICAM-1 expression (from flow cytometry data in panel A) in full growth medium from 3 independent experiments performed in duplicate. Cells are labelled as endothelial cells (ECs), tumour cells (TCs), fibroblasts (HDF) or PBMC. **D)** Cells (grown in full growth media) as in *A*), were lysed and analysed by immunoblotting for the indicated proteins. p-ERK1/2 and p-S6 were detected using antibodies specific for the phosphorylated proteins. Confluency of cells is shown as either confluent (C) or sub-confluent (S); HL60 cells and PBMC are cultured in suspension.

#### Supplementary Figure 2

**A)** BOECs were treated with indicated siRNA for 72 h; surface proteins were then biotinylated and lysates purified on Neutravidin agarose and analysed by immunoblotting. Actin was analysed in an aliquot prior to neutravidin capture (Input). **B)** HUVECs were treated with indicated siRNA for 48 h before lysis and analysis by immunoblotting. **C)** PVR protein expression from panel B) was quantified, normalised to tubulin and expressed relative to treatment with the control siRNA; ANOVA from 3 independent experiments where ** $p<0.01$. **D)** PBMCs were Cell Tracker Green (CTG)-labelled and left to adhere to confluent siRNA-treated HUVECs for 60 minutes before washing in PBS, fixing with 4% PFA and staining with CD45-FITC (green) and CD146-APC (red). **E)** Quantitation of PBMC binding to HUVECs.

Cells treated (as in panel C) along with control HUVEC monolayers pre-treated with anti-PVR function blocking antibodies, were incubated with CTG-labelled PBMCs for 30 minutes before quantification of bound PBMCs; Statistical significance was determined by ANOVA where ** p<0.01. **F)** Migration of siRNA-treated HDLECs as determined with Cell Invasion/Migration (CIM) plates using RTCA. **G)** Quantitation of 3 independent migration experiments, as shown in panel F). Significance was determined by ANOVA where * p<0.01 from 3 independent experiments performed in duplicate. **H)** Angiogenic activity using collagen overlay assay; HDLECs were transfected with siRNA (as indicated) for 48 h prior to overlaying with 1 mg/ml Type I collagen for 18-24 h. Cells were fixed, stained with AF488-phalloidin and imaged using InCucyte live cell imager. **I)** Quantitation of collagen overlay assays; Total branch length, total tubule length, number of branches and number of junctions were assessed from (H) in the mock, control and siPVR transfected HDLECs. Significance was determined by ANOVA where * p<0.05 and ** p<0.01 from 4 independent experiments performed in duplicate. **J)** Angiogenic activity using fibroblast co-cultures: siRNA-treated HDLECs were co-cultured on HDFs for 9 days before fixation and staining with anti-CD31-AF488.

**Supplementary Figure 3.**

**A)** BOECs were transduced with a pool of 3 shRNA-expressing lentiviruses targeting PVR. Clones were selected in puromycin and pooled populations analysed by immunoblotting. Blots are representative of 3 independent transductions. **B)** Quantification of p53 protein expression induced by siRNA against PVR, normalised against actin and expressed relative to siScr levels. Significance was determined by ANOVA where *** p<0.001. **C)** HUVECs treated with various siRNA targeting PVR and/or p53. 72 h post-transfection cells were labelled with anti-PVR antibody before fixation and analysed by immunofluorescent microscopy for PVR, F-actin and p21 expression. Scale bar, 20 μM. **D)** HDLECs treated with indicated siRNA for 24 h prior to culturing onto confluent HDFs for 7d before fixation, staining with anti-CD31 and imaging using InCucyte Live Cell Imager or Operetta HTS. Quantification of images are shown in Fig. 2*G*. **E)** HDLECs treated with various siRNA for 24 h, were grown as hanging drops (to promote spheroid formation) at $3x10^3$ cells/30 μL media for 24 h prior to collection and embedding in 1 mg/mL type I collagen. Cells were overlaid with full EC media and cultured for a further 24 h prior to fixation, staining with phalloidin-AF488, and imaging on an EVOS microscope. Scale bar, 20 μM. **F)** Morphology of siRNA-treated HDLECs. Cells were transfected with the indicated siRNA for 48 h before reseeding onto culture dishes in the absence of an ECM for a further 48 h before examining morphology by phase-contrast microscopy. Images are representative of 2 independent experiments.

**Supplementary Figure 4.**

**A)** hTERT-immortalised HDFs were transduced with control (pLV-Scr) or V5-tagged UL141 (pLV-UL141) and selected with puromycin. Pooled populations were assessed for plasma membrane expression of indicated proteins using cell surface biotinylation followed by Neutravidin agarose pulldown and Western blotting. **B)** HMBECs and BOECs were transduced as in panel A and analysed for changes in surface expression as outlined above. **C)** Co-culture angiogenesis assays of HUVECs expressing UL141 (or control) and HDFs, cultured for 9 days before being fixed and stained with anti-CD31 antibody and analysis of tubule network formation. Network parameters are quantitated in D. **D)** Tubular networks formed by HUVECs expressing UL141 or control cells (as shown in panel C). The data shows network branches, network length, network area, average network length and average tubule width. Significance was determined by one-tailed student t-test from 3 independent experiments where $*p<0.05$. **E)** UL141 (or control) transduced HUVECs treated with doxorubicin (Dox; 0, 0.2μM and 1μM) were assessed for p21, MDM2 and p53 expression by immunofluorescence microscopy (scale bar 50 μm). Quantitation of p53 levels are shown in panel G. **F)** Nuclear p53 levels quantified from 3 independent experiments performed in duplicate and analysed by ANOVA where $*\ p<0.05$ and $**\ p<0.01$**.** Data are expressed relative to the control transduced cells. **G)** Cell viability (using an MTT assay) of UL141 expressing HUVECs (and controls), produced as in A, treated with the topoisomerase II inhibitor, doxorubicin (Dox) for 24 h. Viability is expressed as a percentage relative to the control treated cells and is quantified from 3 independent experiments performed in triplicate using ANOVA and Tukey's post-test where **p<0.01.

### Supplementary Figure 1

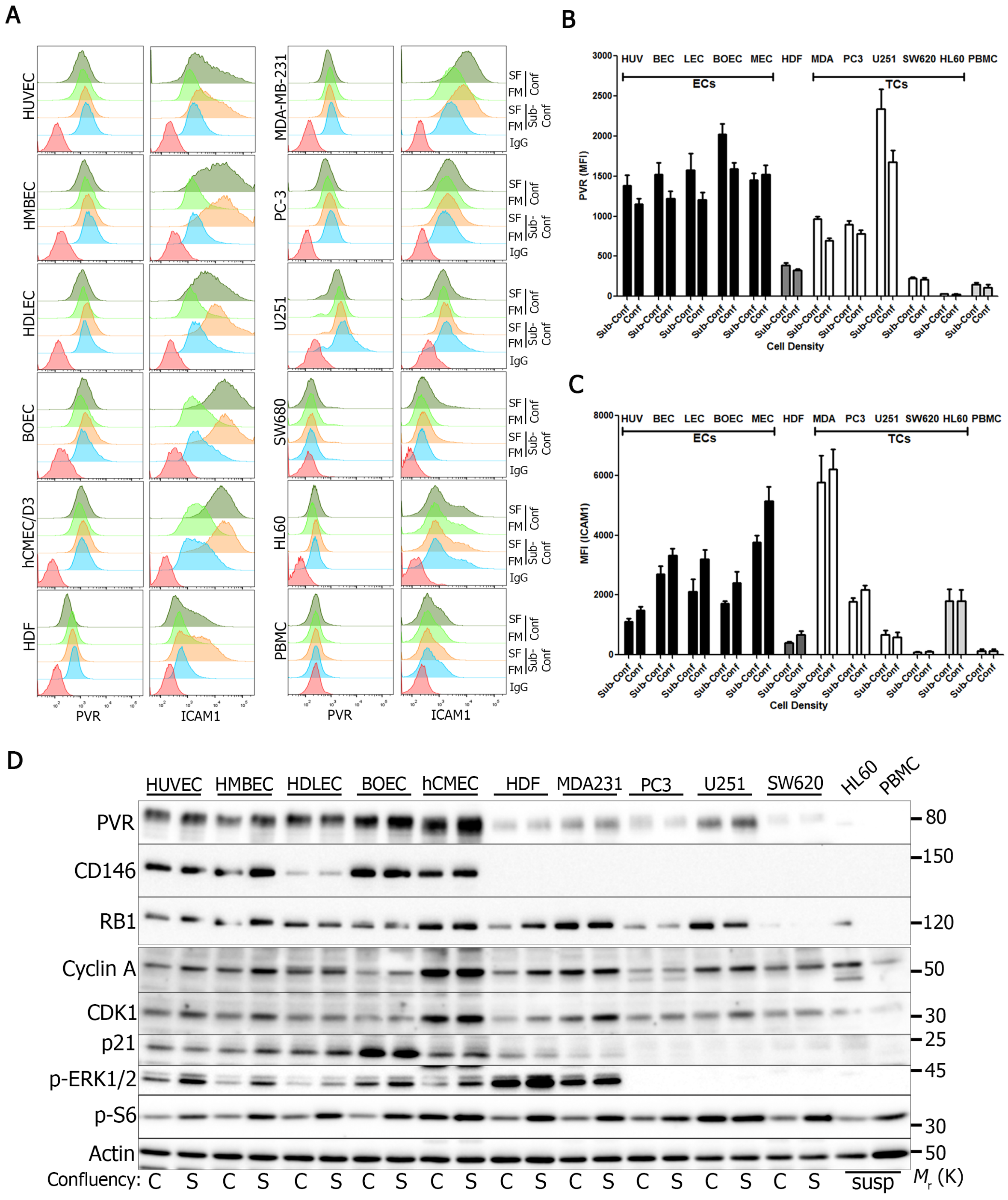

### Supplementary Figure 2

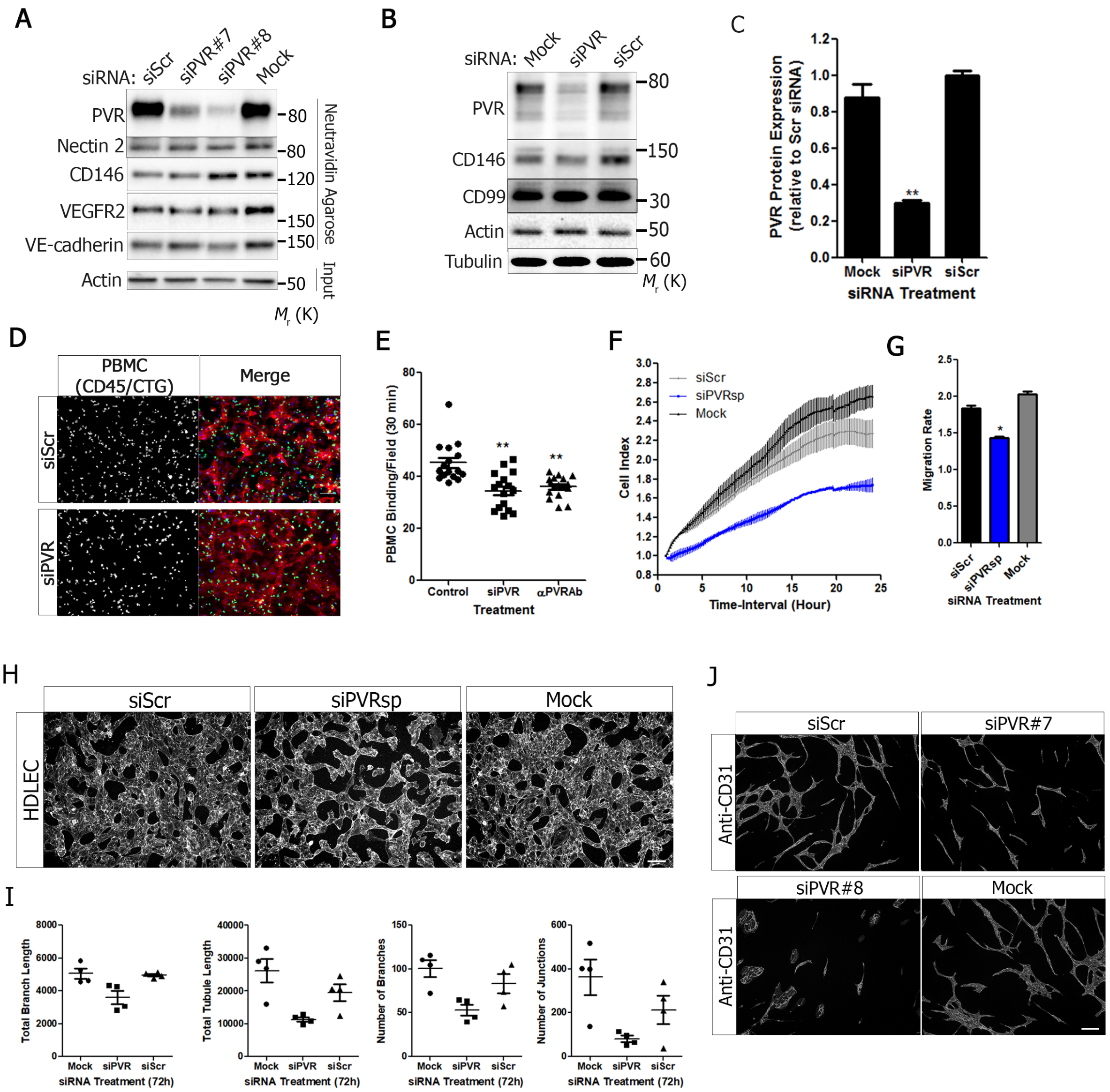

### Supplementary Figure 3

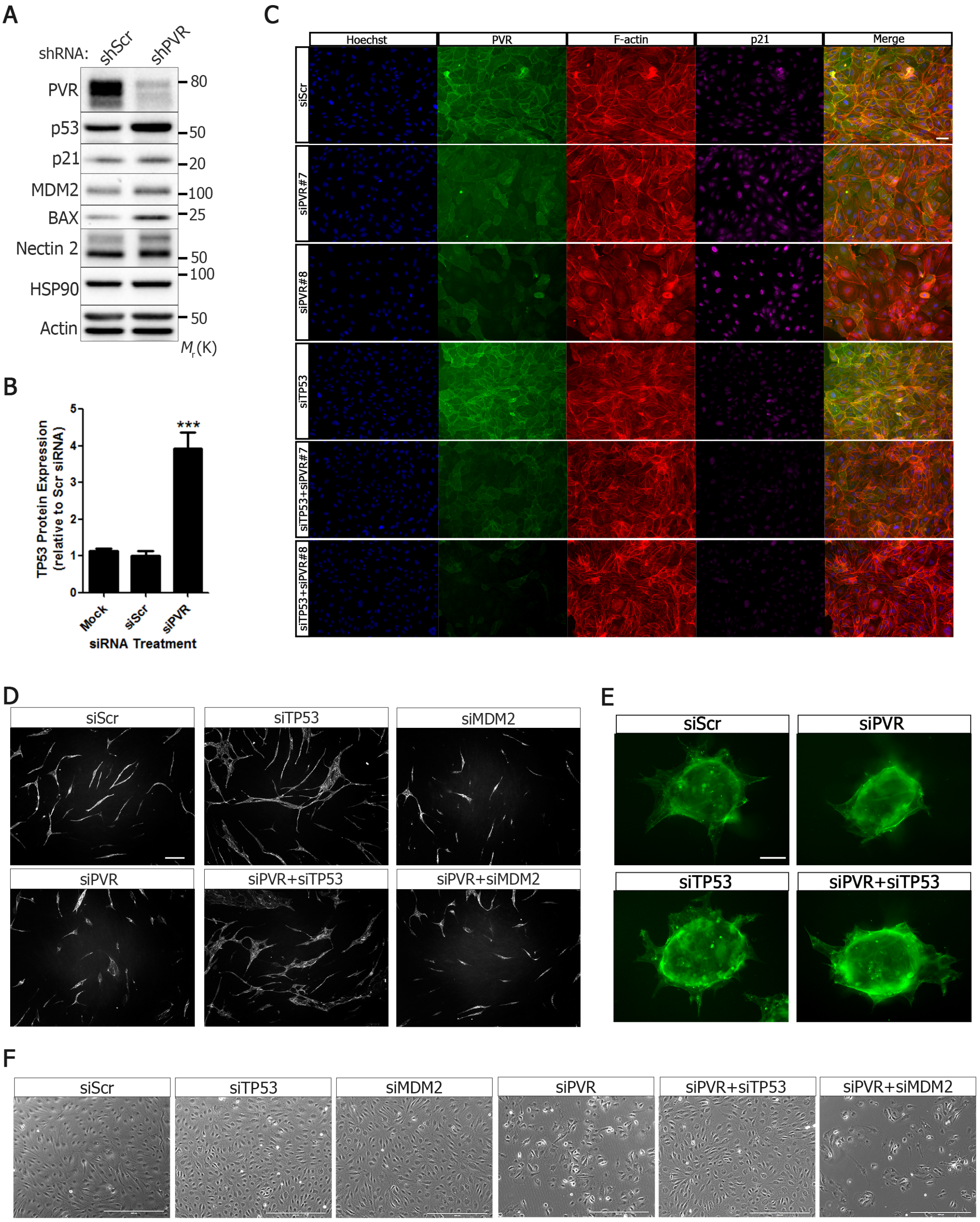

### Supplementary Figure 4

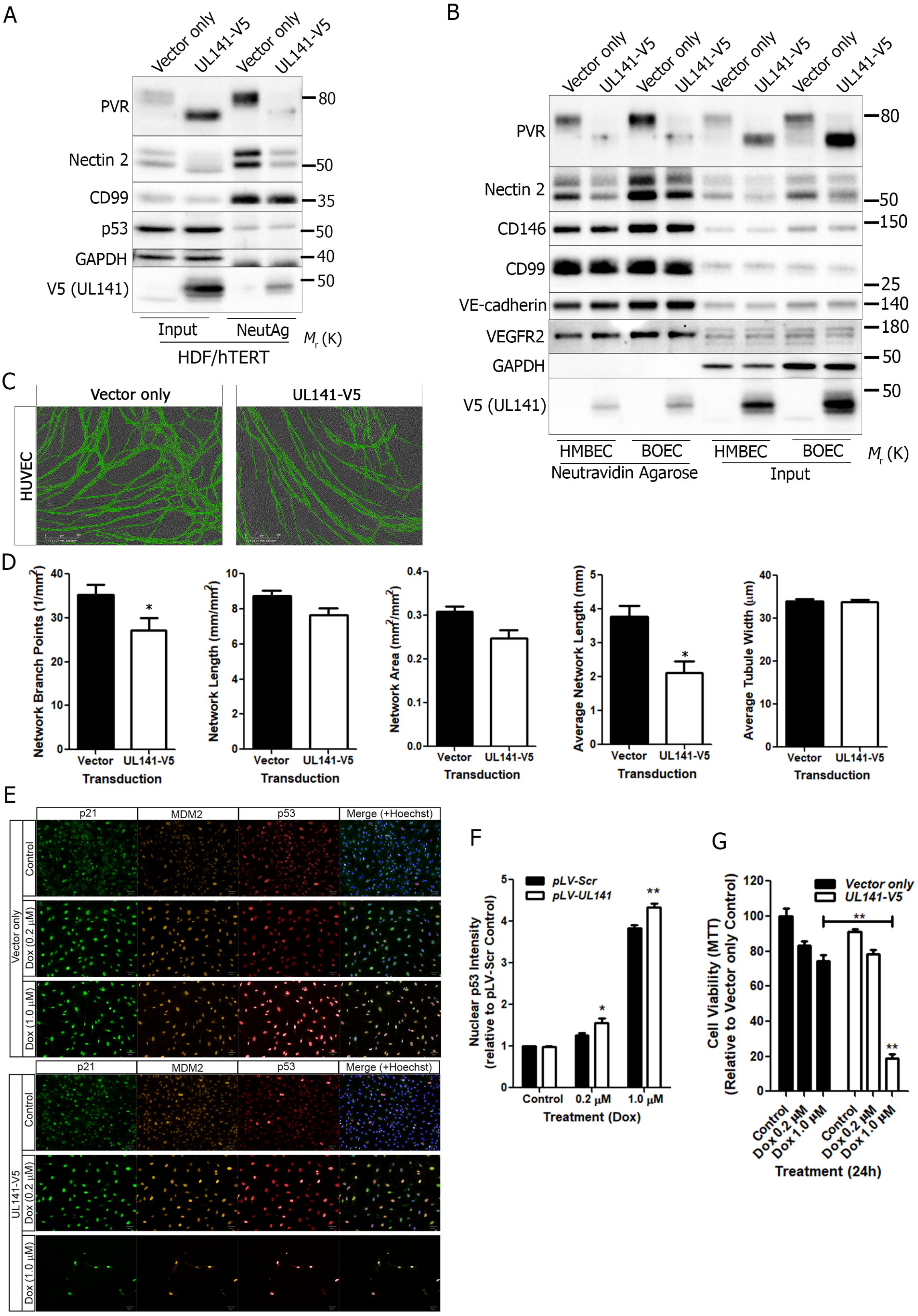
